# Activity-dependent extracellular proteolytic cascade remodels ECM to promote structural plasticity

**DOI:** 10.1101/2024.10.28.620597

**Authors:** Jeet Bahadur Singh, Bartomeu Perelló-Amorós, Jenny Schneeberg, Constanze I. Seidenbecher, Anna Fejtová, Alexander Dityatev, Renato Frischknecht

**Affiliations:** Leibniz Institute for Neurobiology (LIN), 39118 Magdeburg, Germany; Department of Psychiatry and Psychotherapy, University Hospital Erlangen, Friedrich-Alexander-Universität Erlangen-Nürnberg, Erlangen, Germany; Center for Cellular and Molecular Therapeutics (CCMT), Colket Translational Research Building, Children’s Hospital of Philadelphia (CHOP), 3501 Civic Center Blvd, Philadelphia, PA 19104, USA; Department of Biology, Animal Physiology, Friedrich-Alexander-Universität Erlangen-Nürnberg, Erlangen, Germany; German Center for Neurodegenerative Diseases (DZNE), Molecular Neuroplasticity Group, 39120 Magdeburg; Center for Behavioral Brain Sciences (CBBS), 39106 Magdeburg, Germany; Medical Faculty, Otto von Guericke University, 39120 Magdeburg, Germany

## Abstract

The brain’s perineuronal extracellular matrix (ECM) is a crucial factor in maintaining the stability of mature brain circuitry. However, how is activity-induced synaptic plasticity achieved in the adult brain with a dense ECM? We hypothesized that neuronal activity induces cleavage of ECM components, creating space for synaptic rearrangements. To test this hypothesis, we investigated neuronal activity-dependent proteolytic cleavage of brevican, a prototypical perineuronal ECM proteoglycan, and its importance of this process for functional and structural synaptic plasticity in the rat hippocampus *ex vivo*. Our findings revealed that chemical long-term potentiation (cLTP) triggers a rapid brevican cleavage through the activation of an extracellular proteolytic cascade involving proprotein convertases and ADAMTS-4 and ADAMTS-5. This process is dependent on NMDA receptors and requires astrocytes. Interestingly, the extracellular full-length brevican increases upon cLTP, indicating a simultaneous secretion of ECM components. Interfering with cLTP-induced brevican cleavage did not impact the early LTP but prevented formation of new dendritic protrusions. Collectively, these results reveal a mechanism of activity-dependent ECM remodeling and suggest that ECM degradation is essential for structural synaptic plasticity.

## Introduction

A challenge for the adult brain is to create new memories while maintaining stable storage of previously learned information. Memory storage relies on the persistence of these networks, whereas learning and memory formation require functional and structural rearrangements of neuronal networks (Caroni *et al*, 2012). Dendritic spines, the primary sites of excitatory transmission, undergo activity-induced dynamic structural changes that functionally modify neuronal circuits (Bernardinelli *et al*, 2014; Engert & Bonhoeffer, 1999; Holtmaat & Svoboda, 2009). For example, new spines are formed in acute hippocampal slices shortly after the induction of long-term potentiation (LTP). These spines are believed to be formed from dynamic filopodia-like dendritic processes, which are abundant in young animals and persist to a lesser extent in adults (Engert & Bonhoeffer, 1999; Knott *et al*, 2006; Maletic-Savatic *et al*, 1999; Yuste & Bonhoeffer, 2004). In the adult brain, neurons are encased in a specialized extracellular matrix (ECM) that stabilizes synaptic contacts. This may contribute to the limited plasticity observed in adult neuronal networks. Injections of enzymes that degrade the perineuronal ECM into the visual cortex resulted in the restoration of juvenile forms of plasticity (Pizzorusso *et al*, 2002). Moreover, these injections were found to enhance spine motility, indicating that the perineuronal ECM may restrict the morphological alterations of dendritic spines in the visual cortex of adult mice (de Vivo *et al*, 2013).

The perineuronal ECM has two main forms: a condensed form called the perineuronal net (PNN), which is primarily present around a specific group of inhibitory neurons, and a loose form that surrounds almost all neurons and their synapses in the brain (Celio & Blumcke, 1994; Fawcett *et al*, 2022; Lupori *et al*, 2023). Although the two forms of perineuronal ECM differ in their appearance, they are very similar in molecular composition. Both forms consist of a meshwork of glycoproteins and proteoglycans assembled around the glycosaminoglycan hyaluronic acid (Gundelfinger *et al*, 2010). They are rich in chondroitin sulfate proteoglycans (CSPGs), such as aggrecan and brevican of the lectican family, which play a critical role in reducing neuronal plasticity in the adult brain (Fawcett *et al*., 2022; Gundelfinger *et al*., 2010; Sorg *et al*, 2016). Brevican is the most abundant lectican in the mature brain and is found throughout the perineuronal ECM. It is composed of an N-terminal globular domain that binds to hyaluronic acid, a central portion with chondroitin sulfate (CS) attachment sites, and a C-terminal domain that interacts with other ECM molecules or adhesion molecules on the cell surface (Frischknecht & Seidenbecher, 2012; Hedstrom *et al*, 2007).

Brevican and other lecticans, such as aggrecan, are susceptible to proteolytic cleavage by metalloproteases of the ADAMTS family (A Disintegrin and Metalloproteinase with Thrombospondin motifs) family (Nakamura *et al*, 2000). Proteolytic cleavage occurs at the central unstructured region of the protein, separating the N-terminal hyaluronic acid binding domain from the rest of the molecule, thereby loosening the ECM structure (Zimmermann & Dours-Zimmermann, 2008). Here, we hypothesize that this process may represent an endogenous mechanism to disperse the ECM allowing for structural plasticity in the adult brain to promote neuronal network rearrangements and learning.

To test this hypothesis, chemical LTP (cLTP) was induced in acute hippocampal slices of young adult rats, and brevican proteolytic cleavage was quantified using Western blotting. A marked increase in brevican cleavage was observed shortly after cLTP induction, which was abolished by ADAMTS-4/-5 protease inhibitors. Additionally, brevican cleavage at the cleavage site specific for ADAMTS-4/5 required the activity of proprotein convertases and depended on Ca^2+^ signaling through NMDA receptors and voltage-gated calcium channels. Prevention of brevican processing reduced cLTP-induced dendritic spine formation without affecting NMDA-dependent LTP. This suggests that activity-induced remodeling of the perisynaptic ECM was necessary for structural but not functional plasticity in the hippocampus. Thus, neuronal activity triggers a proteolytic cascade that leads to degradation of specific components of the perisynaptic ECM and may open a time window for synaptic rearrangements necessary for learning and memory formation.

## Results

### cLTP promotes the proteolytic cleavage of brevican

To investigate a potential activity-dependent cleavage of the abundant perineuronal ECM proteoglycan brevican, acute hippocampal slices from rats aged 8-12 weeks were used. At this age the mature ECM is fully established. Chemical induction of NMDA receptor-dependent long-term potentiation (cLTP) was achieved by incubating slices for 15 minutes with picrotoxin, forskolin, and rolipram (PFR, Figure 1, Otmakhov *et al*, 2004). Following PFR treatment, the slices were incubated in ACSF for 30 minutes. The ECM was extracted using chondroitinase ABC, a glycosidase enzyme that digests chondroitin sulfate. This procedure releases ECM proteins into the supernatant, facilitating the separation of extracellular proteins from cellular components by centrifugation (Supplementary Figure 1; Deepa *et al*, 2006; Niekisch *et al*, 2019). To evaluate the secretion and cleavage of brevican, we quantitatively analyzed Western blots of supernatants using an antibody directed against the N-terminus of brevican. This antibody detects both the full-length protein of 145 kDa and N-terminal proteolytic fragments of approx. 53 kDa (Ms α BC; Figure 1B, C). At 30 minutes after the PFR stimulation, there was a 28% increase in the 53 kDa proteolytic fragment (Figure 1C, D red box, and see supplementary Figure 2 for a time-course). Similarly, the abundance of the full-length 145 kDa protein increased by 23% after PFR treatment (Figure 1C, E red box). Brevican undergoes proteolytic cleavage between the amino acids Glu^395^-Ser^396^ by members of the ADAMTS metalloprotease family (Nakamura *et al*., 2000). Antibodies directed against the resulting C-terminal neo-epitope (neo) specifically recognize the 53 kDa ADAMTS-derived proteolytic fragment (Figure 1B; Supplementary Figure 3; Matthews *et al*, 2000; Valenzuela *et al*, 2014). The neo antibody detected a 53 kDa band, and quantification revealed a 45% increase in the 53 kDa neo fragment after PFR treatment (Figure 1C, F red box). This indicates that metalloproteases of the ADAMTS family catalyze proteolysis of brevican in acute hippocampal slices after cLTP induction.

**Figure 1:**
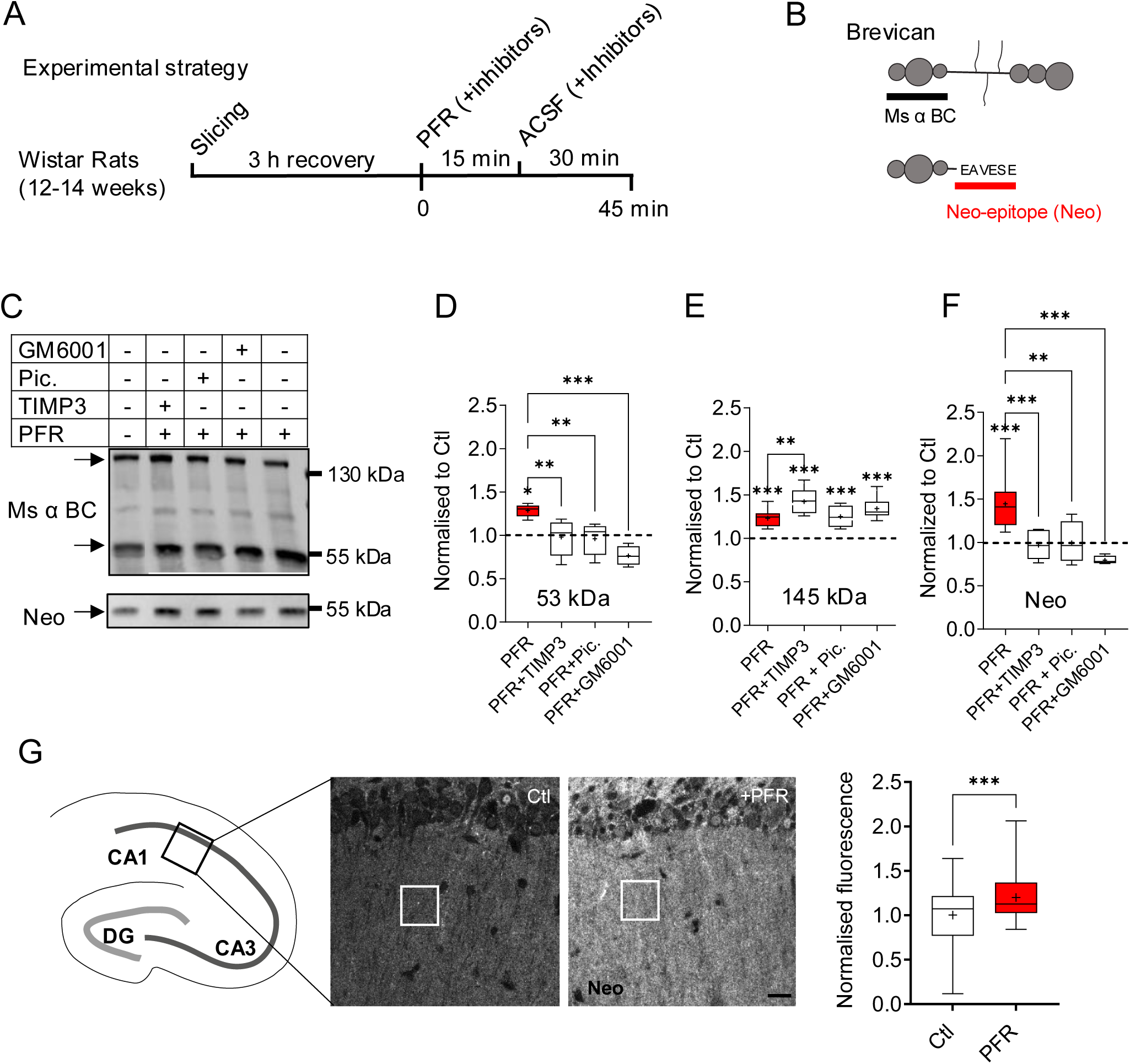
Brevican is proteoytically cleaved in an activity-dependent manner. **A**) Schematic representation of the experimental strategy. **B**) Binding sites of brevican antibodies (black line: Ms anti brevican; red line: rb anti neo epitope). **C**) Representative Western blots of ECM extracts from control and differentially treated hippocampal slices probed with mouse anti-brevican antibody (top) and neo antibody (bottom). Arrows indicate the 145 kDa full-length protein and the cleaved 53 kDa fragment that were quantified in D-F. **D-F**) Quantification of the 53 kDa and 145 kDa bands detected by ms α brevican and neo antibody. Note the significant increase of the 53 kDa fragment after PFA treatment detected by ms anti brevican (D) and neo (F) antibody. The administration of the protease inhibitors TIMP3, piceatannol (pic.) and GM6001 abolished the effect of PFR. **E)** The intensity of the full length 145 kDa protein was significantly increased after PFR treatment. This increase was significantly higher in the PFR-TIMP3 group compared to PFR alone. All data were normalized to protein loading and control slices (dashed line). (Ms α BC, D) 53 kDa: PFR: 1.29 ± 0.03 n = 6; PFR+TIMP3: 0.98 ± 0.07 n = 8; PFR+Pic.: 0.96 ± 0.07 n = 5; PFR+GM6001: 0.77 ± 0.06, n = 4. Mean ± SEM. E) 145 kDa: PFR = 1.23 ± 0.03 n = 13, PFR+TIMP3: 1.42 ± 0.07 n = 5; PFR+Pic.: 1.25 ± 0.06 n = 4; PFR+GM6001: 1.34 ± 0.04 n = 8. F) Neo: PFR = 1.45 ± 0.11 n = 9; PFR+TIMP3: 0.97 ± 0.07 n = 6; PFR+Pic.: 0.99 ± 0.12 n = 4; PFR+GM6001: 0.80 ± 0.02 n = 4; Values compared to Ctl and PFR. One Way ANOVA, Šídák’s multiple comparisons test, * P < 0.05, ** P < 0.01, *** P < 0.001). **G**) Control and PFR-treated acute hippocampal slices were subjected to immunofluorescence staining with neo epitope antibodies. The fluorescence intensity in the neuropil of the CA1 region was compared (boxed area; scale bar, 20 μm). A significant increase in neo staining was observed following PFR treatment within the neuropil (Ctl: 1.0 ± 0.04 n = 71, PFR: 1.20 ± 0.03 n = 63. Unpaired t test, *** P < 0.001). For detailed statistics see Supplementary table.

### Inhibition of ADAMTS prevents brevican processing

To verify the role of ADAMTS proteases in the activity-dependent proteolysis of brevican, we induced cLTP in acute hippocampal slices in the presence of broad-spectrum matrix metalloprotease inhibitors and ADAMTS protease-specific inhibitors (Figure 1A-F). Brevican cleavage was again measured by quantifying the 53 kDa proteolytic fragments detected by the neo and ms anti-brevican antibody on Western blots. cLTP-induced formation of the 53 kDa brevican fragment was completely prevented in the presence of the general metalloprotease inhibitor GM6001 (Figure 1C, D). Similarly, TIMP3, an inhibitor of ADAMTS-4 and −5, and piceatannol, an ADAMTS-4 inhibitor (Lauer-Fields *et al*, 2008), reduced brevican cleavage (Figure 1C, D and F). In addition, a significant increase in the full-length brevican protein was observed during cLTP induction when used with the aforementioned protease inhibitors (Figure 1E). The effects on both the full-length and the 53 kDa fragment of brevican were unaffected by the application of the protein synthesis inhibitor anisomycin during cLTP induction (Supplementary Figure 4A). This indicates that the observed effects are more likely due to changes in brevican secretion and cleavage rather than activity-dependent regulation of brevican synthesis.

### Proteolysis of brevican occurs in synaptic layers of the CA1

Previous publications have demonstrated perisynaptic proteolysis of brevican in dissociated neuronal cultures (Mitlohner *et al*, 2020; Valenzuela *et al*., 2014). Consequently, the aim was to ascertain whether brevican undergoes cleavage in the synapse-rich *stratum radiatum* of the CA1 region of acute hippocampal slices following the induction of cLTP. To achieve this, immunohistochemistry was conducted using the neo antibody to identify the ADAMTS-derived N-terminal fragment of brevican in control and PFR-treated acute hippocampal slices. The proteolytic fragment of brevican was observed throughout the *stratum radiatum* of the CA1 region, as previously reported (Figure 1G; Valenzuela *et al*., 2014; Yuan *et al*, 2002). We quantified fluorescence intensity in the CA1 region and compared PFR-treated slices to control slices (Figure 1G; boxed area). We observed elevated fluorescence intensities after PFR treatment, indicating cleavage of brevican within the loose ECM present in the molecular layer of the CA1 (Figure 1G). Together, these experiments indicate that brevican was cleaved by members of the ADAMTS family in an activity-dependent manner within synaptic regions.

### Brevican cleavage depends on the activity of proprotein convertases with furin-like specificity

Proteases from the ADAMTS family are first produced as inactive zymogens. Their activation relies on the removal of their prodomain by members of the proprotein convertase subtilisin/kexin type (PCSK) protease family, which have a substrate specificity similar to that of furin (Kelwick *et al*, 2015). To evaluate the impact of PCSK on the proteolysis of brevican, three PCSK-specific inhibitors were administered during the induction of cLTP (Figure 2A). All three PCSK inhibitors used in the experiments – Furin inhibitor I (Furini I), Furin inhibitor II (Furini II), and Proprotein Convertase Inhibitor (PCi) - inhibited the activity-dependent formation of the 53 kDa proteolytic fragment of brevican (Figure 2A, B). This suggests that the cLTP-induced cleavage of brevican is dependent on proteolytic cascades involving PCSK-dependent ADAMTS activation.

**Figure 2.**
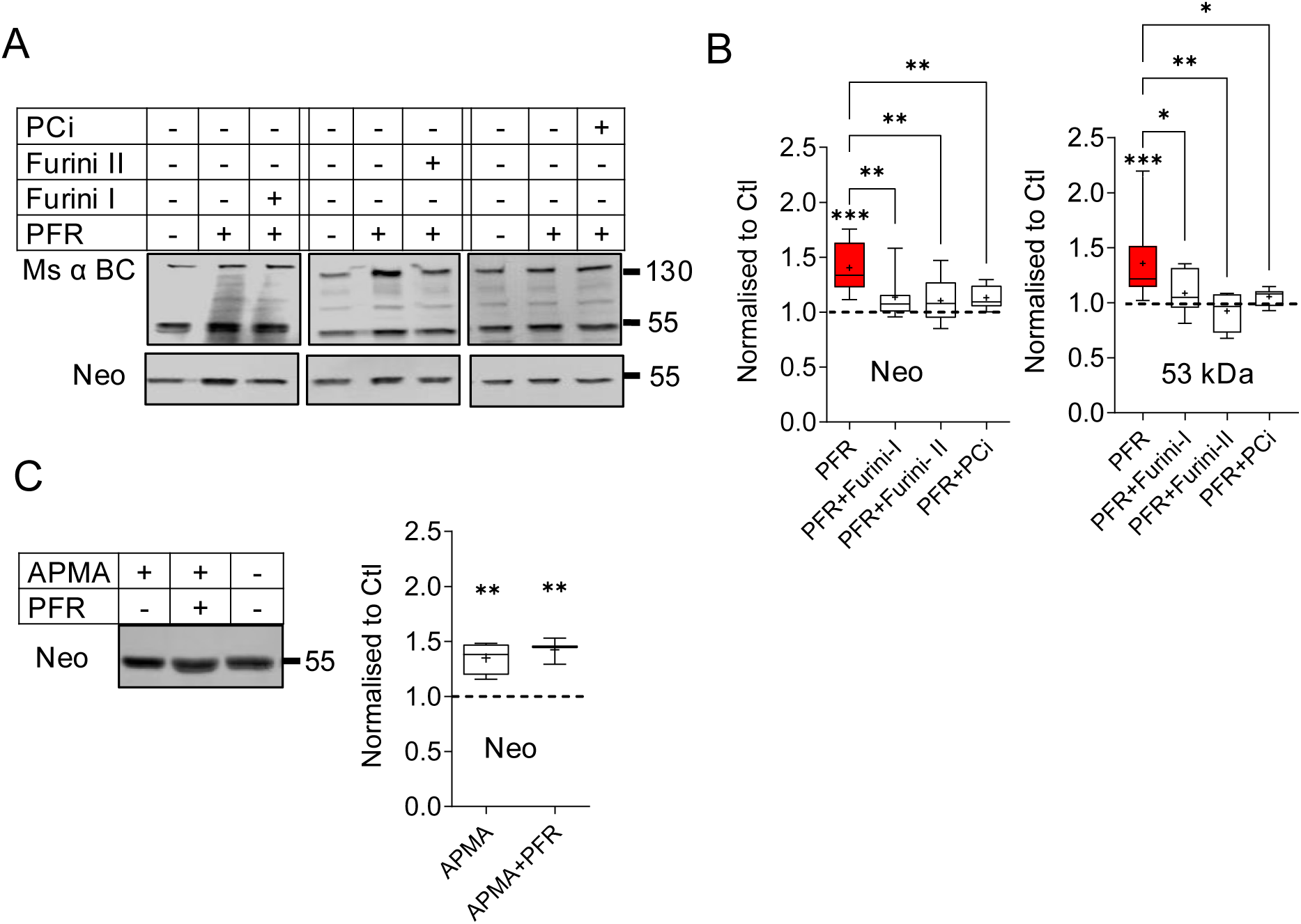
Activity of proprotein convertases is necessary to induce brevican cleavage. **A**) Representative Western blot of extracts from acute hippocampal slices treated with PFR (15 min PFR + 30 min ASFS) in combination with three different inhibitors of PCSK with furin-like specificity (Furini I, Furini II and PCi). **B**) Quantification of the 53 kDa proteolytic band of brevican, detected with ms α brevican or neo antibody. All three PCSK inhibitors abolished PFR-induced brevican cleavage (53 kDa: PFR: 1.36 ± 0.07 n= 21; PFR+Furini I: 1.09 ± 0.06 n = 10; PFR+Furini II: 0.92 ± 0.09 n = 4; PFR+PCi: 1.06 ± 0.03 n = 7; Neo: PFR: 1.40 ± 0.06 n= 14; PFR+Furini I: 1.14 ± 0.06 n = 13; PFR+Furini II: 1.10 ± 0.07 n = 8; PFR+PCi: 1.13 ± 0.04 n = 7. Values compared to Ctl and PFR, mean ± SEM. One Way ANOVA, Šídák’s multiple comparisons test, * P < 0.05, ** P < 0.01, *** P < 0.001). **C**) Western blot and quantification of extracts from acute hippocampal slices treated with APMA and combined treatment with PFR and APMA. Note the significant increase of brevican cleavage under both conditions. All data were normalized to protein loading and control slices (dashed line). (Neo: APMA: 1.35 ± 0.07 n = 4; APMA+PFR: 1.43 ± 0.07 n = 4. Values compared to Ctl indicated by the dashed line, mean ± SEM. One Way ANOVA, Dunnett’s multiple comparisons test, ** P < 0.01). For detailed statistics see Supplementary table.

The PCSK family members can activate their substrate both intracellularly in the secretory pathway and extracellularly (Seidah & Prat, 2012). To determine whether ADAMTS activation occurred intra or extracellularly, slices were incubated with 4-aminophenylmercuric acetate (APMA), a compound that directly activates metalloproteases without cleavage of the prodomain (Peixoto *et al*, 2012; Van Wart & Birkedal-Hansen, 1990). Hippocampal slices treated with APMA resulted in brevican cleavage, even without cLTP induction, as shown in Figure 2C. The combined application of APMA and PFR on slices led to a slight increase in brevican cleavage, although it was not statistically significant, as indicated in Figure 2C. This result suggests the presence of inactive ADAMTS in the extracellular space, with ADAMTS activation being necessary and sufficient to initiate brevican cleavage.

### Brevican cleavage depends on D-serine

Astrocytes play a critical role in synaptic plasticity and regulate several ECM proteins, including brevican (Favuzzi *et al*, 2017; Hamel *et al*, 2005; John *et al*, 2006). To investigate their involvement in brevican processing during cLTP, we administered the gap junction inhibitors carbenoxolone (CBX) and endothelin (endo), which are known to disrupt astrocyte function (Beltran-Castillo *et al*, 2017; Blomstrand *et al*, 1999). The results in Figure 3A show that cLTP induction had no effect on brevican cleavage in the presence of CBX and endo indicating that normal astrocyte function is required for activity-induced brevican cleavage (Figure 3A). Astrocytes support synapse function by promoting the release of D-serine, which acts as a co-agonist of NMDA receptors (NMDARs; Henneberger *et al*, 2010; Panatier *et al*, 2006; Wolosker *et al*, 2016). Indeed, addition of D-serine to slices completely reversed the impact of CBX on brevican cleavage. Moreover, D-serine induced brevican cleavage under basal conditions, i.e., without cLTP induction (Figure 3B). Application of D-serine has been reported to activate postsynaptic NMDA receptors in conjunction with basal neuronal activity (Henneberger *et al*., 2010). Therefore, we investigated the dependence of cLTP-induced brevican cleavage on synaptic glutamate receptor activation. The cleavage of brevican induced by D-serine was abolished by the NMDAR antagonist APV, confirming the requirement of NMDAR signaling in the process (Figure 3B). These findings suggest that astrocytes are involved in the cleavage of brevican triggered by neuronal activity. This is most likely due to their role in providing the necessary D-serine for the activation of NMDARs.

**Figure 3.**
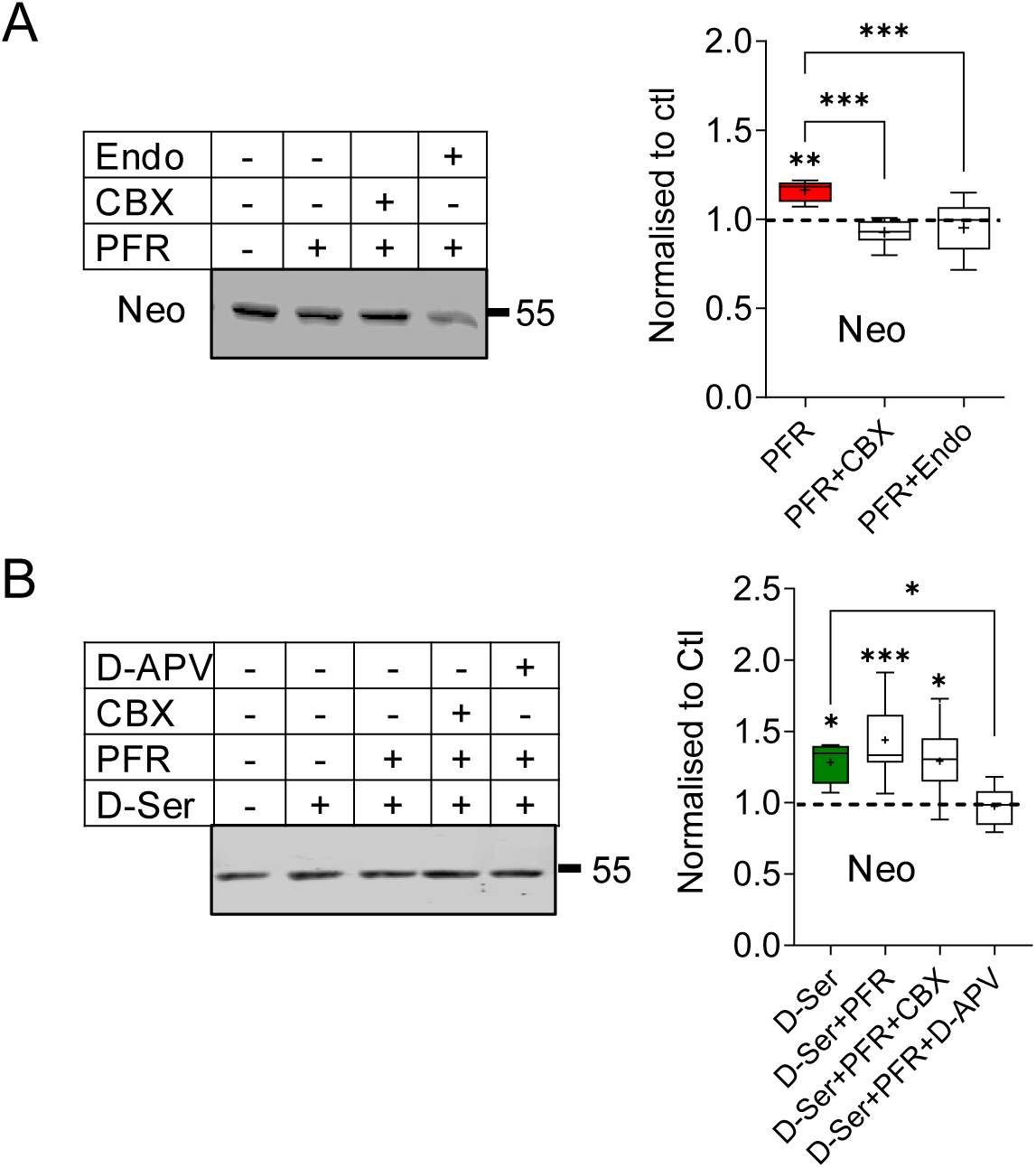
Astroglia affect brevican proteolysis indirectly. **A**) Western blot and quantification of extracts from acute hippocampal slices treated with CBX and endothelin (endo) to interfere with glia function. PFR-induced augmentation of brevican cleavage was abolished by endo and CBX. All values are expressed as a percentage of the control, which is indicated by the dashed line (Neo: 1.17 ± 0,02 n = 7, PFR+CBX: 0.93 ± 0.03 n = 6, PFR+Endo: 0.95 ± 0.06 n = 7. Values compared to Ctl and PFR, mean ± SEM. * P < 0.05, ** P < 0.01, *** P < 0.001). **B**) Western blot analysis and corresponding quantification of slices treated with D-serine, PFR, and the NMDAR blocker APV. D-serine application was sufficient to induce brevican proteolysis. D-serine abolished CBX-induced block of brevican cleavage. In the presence of APV, D-serine did not induce brevican cleavage (Neo: D-Ser: 1.28 ± 0.05 n = 7, D-Ser+PFR: 1.44 ± 0.09 n = 9, D-Ser+PFR+CBX: 1.29 ± 0,08 n = 9, D-Ser+PFR+D-APV: 0,98 ± 0,06 n = 6. Values compared to Ctl and PFR, mean ± SEM. One Way ANOVA, Šídák’s multiple comparisons test, * P < 0.05, ** P < 0.01, *** P < 0.001). All blots were normalized to protein loading and control slices (dashed line). For detailed statistics see Supplementary table.

### Regulation of brevican secretion and cleavage requires activation of the postsynapse

To further analyze the role of glutamate receptors in activity-dependent brevican regulation, specific receptor antagonists were applied during cLTP induction. CNQX, an AMPA type glutamate receptor blocker, was found to eliminate cLTP-induced brevican processing and activity-induced increase in full-length protein (Figure 4A). Treatment with the activity-dependent NMDAR inhibitor MK801 or the blocker of GRIN2B-containing NMDAR RO25-6981 inhibited cleavage but did not affect extracellular accumulation of the full-length protein (Figure 4A). In addition to NMDARs, L-type voltage-gated calcium channels (L-VGCCs) also initiate Ca^2+^ influx and signaling after postsynaptic depolarization, which is necessary for activity-dependent neuroplasticity (Bayer & Schulman, 2019). To investigate the role of L-VGCCs in brevican regulation, we utilized nifedipine, an L-VGCC blocker, during the induction of cLTP. Our findings demonstrate that nifedipine diminished the extracellular upregulation of both full-length and proteolytic fragments of brevican in the slice extracts, which supports the hypothesis that postsynaptic Ca^2+^ acts as second messenger in this process (Figure 4A).

**Figure 4.**
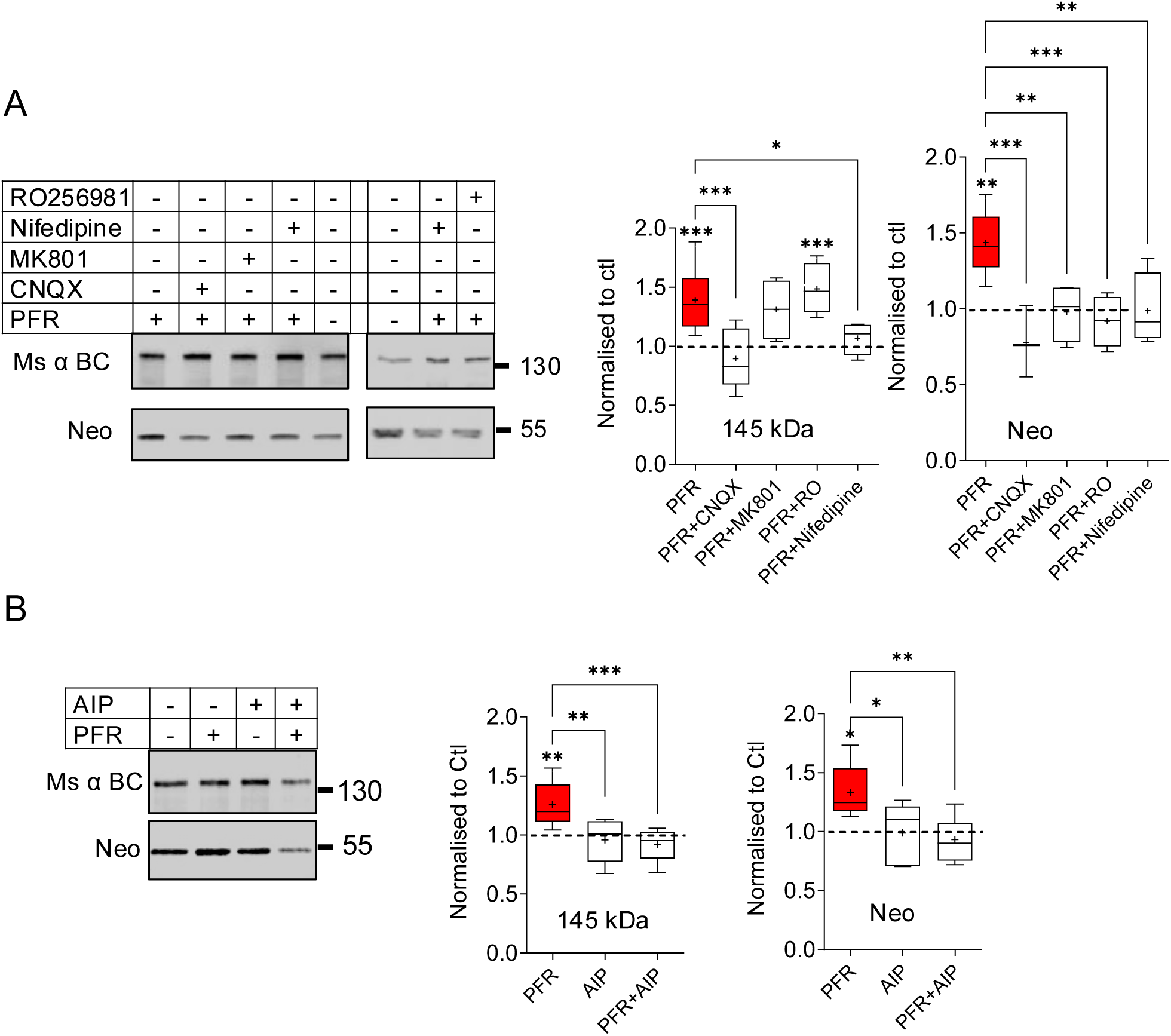
Postsynaptic activation is required for brevican regulation. **A**) Western blot and corresponding quantification of acute hippocampal slices treated with AMPA, NMDA, and L-type calcium channel blocker. The PFR-induced increase in full-length brevican was not observed in the presence of CNQX or nifedipine. The PFR-induced increase of the 53 kDa band detected by the neo antibody is prevented in the presence of CNQX, MK801, RO256981, and nifedipine (145 kDa: PFR: 1.39 ± 0.06 n = 15, PFR+CNQX: 0.90 ± 0.13 n = 5, PFR+MK801: 1.31 ± 0.13 n = 4, PFR+RO: 1.49 ± 0.11 n = 4, PFR+Nifedipine: 1.07 ± 0.07 n = 4. Neo: PFR: 1.44 ± 0.07 n = 9, PFR+CNQX: 0.98 ± 0.10 n = 3, PFR+MK801: 0.98 ± 0.1 n = 4, PFR+RO: 0.92 ± 0.08 n = 4, PFR+Nifedipine: 0.99 ± 0.12 n = 4. Values compared to Ctl and PFR, mean ± SEM. One Way ANOVA, Šídák’s multiple comparisons test, * P < 0.05, ** P < 0.01, *** P < 0.001) **B**) Western blot and corresponding quantification of acute hippocampal slices treated with the CAMKII blocker AIP. PFR-induced increase of the full-length (145 kDa) and cleaved fragment of brevican, as detected by the neo antibody was abolished (145 kDa: PFR: 1.26 ± 0.07 n = 7, AIP: 0.96 ± 0.08 n = 5, PFR+AIP: 0.92 ± 0.05 n = 8. Neo: PFR: 1.33 ± 0.1 n = 5, AIP: 0.99 ± 0.12 n = 5, PFR+AIP: 0.93 ± 0.06 n = 8. Mean ± SEM. Values compared to Ctl and PFR. One Way ANOVA, Šídák’s multiple comparisons test, * P < 0.05, ** P < 0.01, *** P < 0.001). All blots were normalized to protein loading and control slices (dashed line). For detailed statistics see Supplementary table.

To further investigate this hypothesis, we utilized autocamtide-2-related inhibitory peptide (AIP), a specific inhibitor of Ca^2+^/calmodulin-dependent protein kinase II (CaMKII), to block one of the primary plasticity-related Ca^2+^-dependent postsynaptic signaling molecules. AIP was found to diminish the full-length form and proteolytic fragment of brevican, indicating a crucial role of Ca^2+^-signaling in both the cleavage and secretion of brevican upon postsynaptic depolarization (Figure 4B). While the activation of AMPAR, L-VGCC, and CaMKII regulates both the secretion and cleavage of brevican, NMDAR activation appeared to be solely responsible for its proteolysis but not for externalizing full-length brevican.

### Brevican cleavage is not necessary to induce LTP

Next we investigated whether brevican proteolysis by ADAMTS was necessary for LTP induction. As a first approach we quantified in phosphorylation of CaMKII and extracellular signal-regulated kinase (ERK) following PFR treatment on Western blots, two molecular indicators of LTP initiation (Sweatt, 2001; Thomas & Huganir, 2004; Bayer & Schulman, 2019). We blocked ADAMTS activity during cLTP induction utilizing the protease inhibitor TIMP3. However, we found no changes in cLTP-induced phosphorylation of ERK and CamKII (Figure 5A).

**Figure 5.**
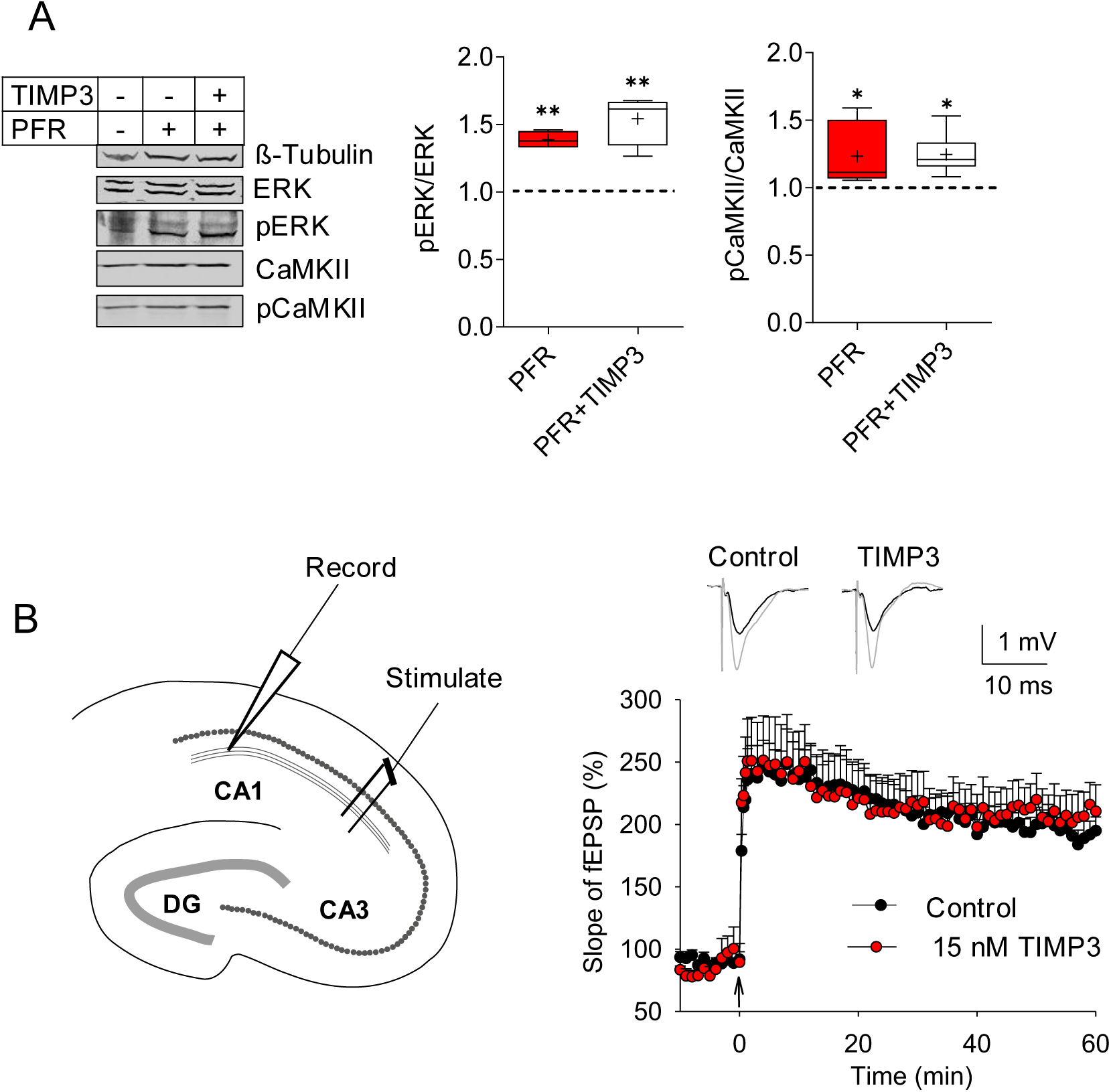
Inhibition of ADAMTS proteases does not affect LTP induction. **A**) Western blots and corresponding quantification of cell lysates from hippocampal slices treated with PFR. The ratio of phosphorylated and unphosphorylated kinases ERK and CaMKII was compared between slices treated with PFR and PFR+TIMP3. The gel loading was adjusted according to the β-tubulin signal. PFR induced ERK and CaMKII phosphorylation, which was not affected by additional TIMP3 treatment (pERK/ERK: PFR: 1.39 ± 0.03 n = 4, PFR+TIMP3: 1.54 ± 0.09 n = 4. pCaMKII/CaMKII: PFR: 1.23 ± 0.08 n = 7, PFR+TIMP3: 1.25 ± 0.06 n = 7. Mean ± SEM. Values compared to Ctl and PFR. One Way ANOVA, Šídák’s multiple comparisons test, * P < 0.05, ** P < 0.01, *** P < 0.001). **B)** Recordings of field excitatory postsynaptic potentials (fEPSPs) in the stratum radiatum of the CA1b were obtained. Stimulation pulses were applied to Schaffer collaterals, and LTP was induced by applying five theta bursts. A robust increase in the slope of fEPSPs was observed in control (black) and in slices treated with TIMP3 (red). For detailed statistics see Supplementary table.

Additionally, we induced early LTP in acute hippocampal slices by theta burst stimulation of the Schaffer collateral to CA3-CA1 pathway. As in the previous experiment, we used the ADAMTS inhibitor TIMP3 to inhibit brevican cleavage during stimulation (Figure 5B). Consistent with the biochemical results, TIMP3 had no effect on the expression of early LTP. This suggests that ADAMTS activity, and consequently, brevican proteolysis, is not essential for LTP initiation (Figure 5B).

### Formation of cLTP-induced dendritic protrusions requires the activity of ADAMTS proteases

LTP leads to the formation of filopodia-like dendritic protrusions, structural changes of existing and formation of new dendritic spines (Engert & Bonhoeffer, 1999; Jourdain *et al*, 2003). In order to ascertain whether brevican proteolysis was necessary for LTP-induced structural plasticity, we induced cLTP using PFR in acute hippocampal slices from mice expressing YFP in sparse neurons, with and without TIMP3. Subsequently, the length and number of dendritic protrusions were measured to enable comparison of the conditions (Figure 6A-C). As previously reported, a significant increase in the number and length of dendritic protrusions was observed in PFR-treated slices compared to control slices (Figure 6C-E; Matsumoto-Miyai *et al*, 2009). However, in the presence of the protease inhibitor TIMP3, the number and size of dendritic protrusions were identical to those in control slices (Figure 6C-E). This indicates that the activity of proteases from the ADAMTS family was necessary for activity-induced structural plasticity of dendritic protrusions in the hippocampus.

**Figure 6.**
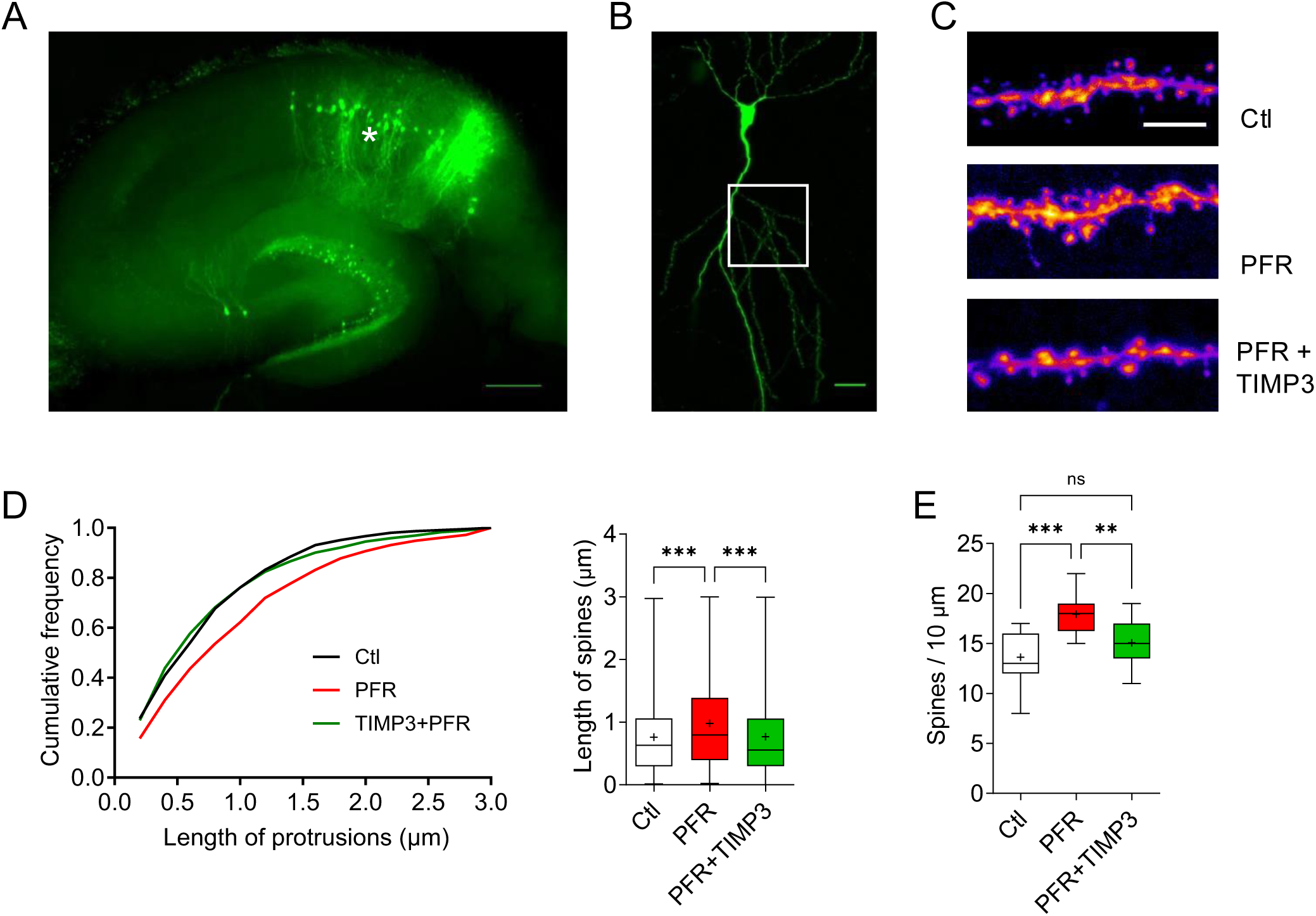
The formation of dendritic spines in response to PFR is inhibited by TIMP3. **A**) Acute hippocampal slice from a mouse expressing YFP in sparse neurons. Asterisk indicates CA1 region (scale bar 500 µm). **B**) Example of YFP expression in CA1 neuron. Spines were counted from the secondary dendrite (Boxed area, scale bar 20 µm). **B)** Examples of dendrites used for quantification. Note the long spines in PFR-treated slices. (arrow, scale bar 5 μm). **D**) Left: Cumulative distribution of dendritic spine length from control (black), PFR (red) and TIMP3+PFR (green) treated slices. Note the shift towards larger spines after PFR treatment (red). Right: Statistical representation of data used for the cumulative representation (Length of spines in µm: Ctl: 0.76 ± 0.03 n = 462, PFR: = 0.98 ± 0.02 n = 1140, PFR+TIMP3: 0.77 ± 0.02 n = 677. Mean ± SEM. Values compared to Ctl and PFR. One Way ANOVA, Šídák’s multiple comparisons test, * P < 0.05, ** P < 0.01, *** P < 0.001, data in supplementary table 6). E) Significant higher number of dendritic protrusion after PFR treatment. This effect is reversed in presence of TIMP3 (Protrusions/10 µm: Ctl: 13.63 ± 0.66 n = 19, PFR: 17.94 ± 0.49 n = 16, PFR+TIMP3: 15.06 ± 0.57 n = 17. One Way ANOVA, Dunnett’s Multiple Comparison Test, * P < 0.05, ** P < 0.01, *** P < 0.001). For detailed statistics see Supplementary table.

## Discussion

The aim of this study was to investigate the activity-dependent remodeling of the perineuronal ECM, the molecular and cellular players involved in this process, and its relevance for functional and structural activity-induced synaptic plasticity. We demonstrate that brevican, an abundant ECM molecule, is proteolytically cleaved during cLTP induction in the synaptic layers of acute hippocampal slices. The processing of brevican is conducted by proteases belonging to the ADAMTS family, which are predominantly activated in the extracellular space by PSCKs. Initiation of this proteolytic cascade depends on NMDA receptors, L-type VDCCs, and CaMKII. Interestingly, inhibiting proteases of the ADAMTS family did not disrupt early LTP. However, it reduced the number of dendritic protrusions typically formed after cLTP induction. Thus, perineuronal ECM modulation appears to occur in an activity-dependent manner and is a prerequisite for structural, but not functional plasticity.

### Neuronal activity induces brevican processing

Activity-dependent extracellular proteolysis and the resulting changes in the ECM have the potential to enhance neuronal plasticity by altering cell adhesion and ECM architecture (Dityatev *et al*, 2010). Indeed, previous studies have demonstrated that ECM proteins and cell adhesion molecules, including agrin and neuroligin-1, undergo proteolytic cleavage in an activity-dependent manner that affects neuronal network architecture and learning (Ferrer-Ferrer *et al*, 2023; Matsumoto-Miyai *et al*., 2009; Peixoto *et al*., 2012). The present study demonstrates that cLTP induces brevican processing, indicating that the perineuronal ECM undergoes acute, activity-dependent remodeling. Our findings indicate that brevican proteolysis relies on NMDAR, L-type Ca2+ channel, and CaMKII activation, which are critical factors in triggering LTP. Our findings further suggest that astrocytes are essential for supporting the expression of NMDA-dependent LTP but do not play a direct role in brevican cleavage, such as through secretion of ADAMTS proteases. In conclusion, these findings indicate that brevican proteolysis is initiated during LTP induction, which may result in ECM remodeling. Furthermore, we observed the proteolytic cleavage of aggrecan, indicating a potential general activity-dependent proteolysis of lectican family members (see Supplementary Figure 4B). Our recent findings further demonstrate that brevican is cleaved under conditions of homeostatic plasticity or after activation of the D1-like dopamine receptor (Mitlohner *et al*., 2020; Valenzuela *et al*., 2014). This suggests that brevican cleavage is induced during different modes of synaptic plasticity and may therefore support these processes.

### Regulation of brevican proteolysis

For local and regulated ECM weakening, activity of proteases must be tightly controlled. This can be achieved by 1) regulating the synthesis of proteases, 2) regulating their activation and/or their 3) externalization from internal stores. Based on our experiments we could exclude protein synthesis of proteases as a critical step since treatment of hippocampal slices with anisomycin, a potent blocker of protein synthesis, in conjunction with PFR did not alter brevican cleavage (see Supplementary Figure 4A).

Proteases of the ADAMTS family are produced as inactive zymogens to prevent their ectopic activity. To activate the enzyme, the pro-domain must be removed from the protein by proteolysis (Kelwick *et al*., 2015). The pro-domains of ADAMTS contain a consensus sequence for pro-protein convertases (PCSK) with furin-like specificity (Kelwick *et al*., 2015). Nine PCSKs have been identified of which seven comprise a furin-like specificity, making them good candidates for ADAMTS activation (Seidah & Prat, 2012). We used three inhibitors of PCSKs with furin-like activity and all three prevented brevican cleavage. This is in line with previous findings showing that PCSKs were necessary to activate ADAMTS in neuronal cultures (Lemarchant *et al*, 2014; Mitlöhner *et al*., 2020; Tortorella *et al*, 2005).

Activation of ADAMTS may occur within the secretory pathway, resulting in secretion of the active enzyme, or extracellularly within the ECM. We incubated acute hippocampal slices with the compound APMA that activates matrix metalloproteases non-proteolytically (Peixoto *et al*., 2012; Van Wart & Birkedal-Hansen, 1990). If brevican was activated within intracellular compartments we expected no additional brevican cleavage upon APMA treatment. However, APMA was sufficient to induce brevican cleavage, suggesting ADAMTS to be localized extracellularly in its inactive form under basal conditions following activity dependent extracellular activation. ADAMTS family members have been reported to bind sulfated glycosaminonglycans on lecticans like aggrecan or brevican (Flannery *et al*, 2002). Thus, our results indicate that activation of ADAMTS to occur extracellularly within the ECM by PCSKs with furin-like activity.

### ECM remodeling is necessary for structural but not functional plasticity

Subsequently, the impact of brevican cleavage on the expression of synaptic plasticity was evaluated. Notably, the blockade of ADAMTS protease activity, which results in brevican cleavage, did not affect the expression of early LTP for 60 min upon induction, as demonstrated by electrophysiological measurements (Figure 5). Nevertheless, our findings indicate that brevican cleavage by ADAMTS proteases is essential for the formation and elongation of dendritic protrusions, which are typically observed following the induction of cLTP (Matsumoto-Miyai *et al*., 2009). This finding is analogous to the results obtained in the neurotrypsin knockout mouse, in which structural plasticity was impaired, but LTP was unaffected (Matsumoto-Miyai *et al*., 2009). The 22 kDa neurotrypsin-derived proteolytic fragment of agrin that is formed during LTP formation has been demonstrated to promote the formation of dendritic protrusions during LTP (Matsumoto-Miyai *et al*., 2009; Stephan *et al*, 2008). It is notable that the N-terminal proteolytic fragment of brevican, but not the full-length protein, has been demonstrated to promote glioma cell motility and invasiveness, suggesting that this fragment may have motility-promoting effects (Hu *et al*, 2008; Viapiano *et al*, 2008). In conclusion, it can be postulated that proteolytic cleavage of brevican produces a fragment, so-called matricryptin, which promotes structural plasticity after LTP induction without affecting LTP expression for at least 60 min.

### Functional impact of ECM degradation

In the mature brain, the perineuronal ECM is a compact molecular network composed of CSPGs and glycoproteins bound to by hyaluronic acid. Studies have shown that enzymatic degradation of the perineuronal ECM by direct injection of glycosidases into the brain is able to restore juvenile modes of plasticity and modify neuronal network activity both *in vitro* and *in vivo* (Bikbaev *et al*, 2015; Carulli *et al*, 2010; El-Tabbal *et al*, 2021; Pizzorusso *et al*., 2002; Rowlands *et al*, 2018). Furthermore, at the cellular level, experimental ECM degradation increased dendritic spine motility in the visual cortex and spine density in the retrosplenial cortex (Cangalaya *et al*, 2024; de Vivo *et al*., 2013). This highlights the restrictive nature of the mature ECM on neuronal plasticity while revealing a significant degree of plasticity in the adult brain. Endogenous mechanisms that alter ECM integrity may therefore have a significant impact on adult brain plasticity and may be necessary for lifelong learning and memory formation.

Supporting, the role of ECM remodeling in learning, the ECM protein brevican is downregulated during the early stages of training in an auditory cortex-dependent learning paradigm (Niekisch *et al*., 2019). Furthermore, exposure to an enriched environment that promotes brain plasticity, also leads to a decrease in brevican expression in animals, highlighting the importance of ECM regulation for neuronal plasticity (Favuzzi *et al*., 2017; van Praag *et al*, 2000). The majority of CSPGs belonging to the lectican family, including brevican and aggrecan, are subject to proteolytic cleavage by metalloproteases of the ADAMTS family, which exhibit aggrecanase-like activity (Kelwick *et al*., 2015; Nakamura *et al*., 2000; Tortorella *et al*, 2002; Zimmermann & Dours-Zimmermann, 2008). Thus, ADAMTS family proteases may remove CSPGs by local activity-dependent extracellular proteolysis, as observed for brevican, allowing higher modes of neuronal plasticity. Recently, the role of microglia has also been highlighted in this context (Nguyen *et al*, 2020; Strackeljan *et al*, 2021; Venturino *et al*, 2021). Microglia were found to engulf ECM proteins and depletion of microglia increased the expression of perineuronal ECM leading to changes in synapse density (Nguyen *et al*., 2020; Strackeljan *et al*., 2021).

In addition to the proteolytic cleavage of brevican, we observed an increase in full-length brevican after cLTP induction (Figure 1C, D). This is enhanced, when ADAMTS or PCSK inhibitors, which prevent brevican processing, are included in the experiment (Figure 2B). Replenishment of the ECM was also observed *in vivo* after enzymatic digestion of the ECM, highlighting the importance of an elaborated ECM for normal adult brain function (Happel *et al*, 2014). Moreover, training mice in an auditory cortex-dependent learning task resulted in a reduction in the abundance of ECM, whereas successful learning led to a marked upregulation of brevican expression, thereby restoring basal levels of brevican (Niekisch *et al*., 2019). It has been proposed that a dynamic focal cluster of perisynaptic ECM, and particularly the sulfation pattern of the chondroitin sulfate side chains of versican, may contribute to activity-dependent plasticity and memory in mice (Chelini *et al*, 2024). It has been demonstrated that components of the ECM are removed by endocytosis and subsequently recycled in an activity-dependent manner (Dankovich *et al*, 2021). Therefore, extracellular proteolysis, microglia, ECM production and recycling may act synergistically *in vivo* to rapidly remove local ECM proteins, thereby creating a window of opportunity for neuronal network readjustment, which is necessary for learning and memory formation. Subsequent replenishment of ECM may be necessary to stabilize newly formed synapses and memories.

## Material & Methods

**Table1.**

| Compound | Supplier | Concentration |
| --- | --- | --- |
| Autocamtide-2 | Sigma-Aldrich | 20 μM |
| Anisomycin | Sigma-Aldrich | 20 μM |
| Carbenoxolone disodium salt (CBX) | Sigma-Aldrich | 100 μM |
| Chondroitinase (ChABC) | Sigma-Aldrich | 0.2 U/ml |
| D-(-)-2-Amino-5-phosphonopentanoic acid (APV) | Tocris Bioscience | 50 μM |
| D-serine | Merck Millipore | 10 μM |
| Furin Inhibitor I | Merck Millipore | 30 μM |
| Furin Inhibitor II (Hexa-D-arginine) | Tocris Bioscience | 0.58 μM |
| Endothelin-1 (endo) | Sigma-Aldrich | 1 μM |
| GM6001 | Tocris Bioscience | 25 μM |
| Picrotoxin | Sigma-Aldrich | 50 μM |
| Nifedipine | Tocris Bioscience | 50 μM |
| 6-Cyano-7-nitroquinoxaline-2,3-dione disodium (CNQX) | Tocris Bioscience | 10 μM |
| Forskolin | Tocris Bioscience | 50 μM |
| Rolipram | Tocris Bioscience | 0.1 μM |
| Ro 25-6981 | Tocris Bioscience | 1 μM |
| TIMP3 | R&D System | 15 nM |

### Acute hippocampal slice preparation

8-12 week old Wistar rats were anesthetized with isoflurane (4%). After decapitation, the brain was rapidly removed and immersed in an oxygenated ice-cold artificial cerebrospinal fluid (ACSF, 125 mM NaCl, 2.5 mM KCl, 1.25 mM NaH2PO4, 25 mM NaHCO3, 2 mM CaCl2, 1 mM MgCl2, 25 mM glucose). Hippocampi were isolated in an oxygenated ice-cold ACSF and transverse hippocampal slices (350 μm) were prepared using a Mcllwain tissue chopper (Mickle Laboratory). The slices were allowed to recover at room temperature for 90 min and then recovered at 32°C for 1 h. Throughout the procedure, the slices were perfused with ACSF solution which was bubbled with 95% O2 and 5% CO2.

### Induction of cLTP in acute hippocampal slices

Chemical LTP (cLTP) was induced with a combination of picrotoxin, forskolin and rolipram (PFR) in ACSF without Mg^2+^ for 15 min at 32°C as previously described (Matsumoto-Miyai *et al*., 2009). All the inhibitors were incubated 20-30 min before stimulation and were present until the end of the recovery after the chemical stimulation (Figure1A, 2A). The list of pharmacological agents and their working concentrations are mentioned in supplementary table Table 1. After treatment, the hippocampal slices were collected in ECM extraction buffer.

### Sample preparation and Quantitative Western blot

For ECM extraction acute hippocampal slices were incubated for 30 min at 37°C in extraction buffer containing chondroitinase ABC (0.2 U/ml) in 0.1 M Tris-HCl, pH 8.0, 0.03 M sodium acetate, complete protease inhibitor cocktail (Roche) and PhosSTOP phosphatase inhibitor cocktail (Roche). For three slices 90 μl of the ECM extraction buffer was used. Slices were triturated and centrifuged at 14000 x g for 10 min. The supernatant was collected and mixed with 4 x SDS Laemmli buffer and stored at −80°C. Protein concentration was determined by the Amido Black colorimetric assay. For SDS-PAGE, gradient gels (tris-glycin 5-20%) containing 2,2,2-trichloroethanol (TCE) were used to visualize and quantify the total amount of protein (Ladner *et al*, 2004). Each fraction was loaded twice and gels were blotted onto polyvinylidene difluoride membranes (Millipore) using the Mighty Small Transfer Tank system (Hoefer). PVDF membranes were blocked in TBS-T containing 0.1% Tween20 and 5% bovine serum albumin for 30 min at room temperature. Primary antibodies (ms anti-BC, 1:1000; BD Transduction Laboratories; Rb anti-neo 1:1000, home-made (Valenzuela *et al*., 2014) were diluted in TBS-T containing 2% BSA and probed for 2 h at room temperature or overnight at 4 °C. Membranes were subsequently incubated with fluorescence coupled secondary antibodies (goat anti-mouse IRDye™-800CW or IRDye™-680RD, goat anti-rabbit IRDye™-800CW or IRDye™-680RD; LI-COR). Blots were scanned using the LI-COR Odyssey system and signals were quantified using ImageJ. Signal intensity was corrected for protein load as determined by TCE for ECM extracts, with GAPDH or β-actin for cell lysates before normalization to the average value of the control group loaded on each membrane.

### Immunocytochemistry

The acute hippocampal slices were fixed overnight in PBS containing 4% PFA and 4% sucrose. They were then placed in 30% sucrose for cryoprotection and frozen in Tissue-Tek (Sakura) embedding medium. The slices were then resliced using a cryostat (Leica CM 3050 S) into 30 μm sections, permeabilized in phosphate-buffered saline (PBS) containing 0.3% Triton X-100 and 10% fetal calf serum (FCS) for one hour at room temperature. The neo-antibody was incubated for 72 hours at 4°C, and the secondary antibodies were incubated overnight at 4°C. Between each staining step, the slices were washed three times for ten minutes in phosphate-buffered saline (PBS). The slices were then mounted on a glass slide using Fluoromount-G™. Images were captured using a Leica SP5 confocal microscope (LAS AF software, version 2.0.2, 63 x objective, NA 1.40, or 20x objective, NA 0.8). For quantitative immunofluorescence, 10 to 12 optical sections were taken with a step size of 250 nm. Maximum intensity projections were generated using the ImageJ software (National Institutes of Health, http://rsb.info.nih.gov/ij/). Fluorescence intensity was quantified within 50 µm rectangular regions within the molecular layer of the CA1 region using the ImageJ software.

### Quantification of dendritic protrusions

For the quantification of dendritic protrusions, acute hippocampal slices from adult Slick V-cre YFP-expressing mice were utilized, as these mice express YFP in sparse neurons. Slices were prepared for imaging as previously described. For the analysis of dendritic protrusions, images of apical secondary dendrites 80 μm from the soma were obtained using a Leica SP5 confocal microscope (LAS AF software, version 2.0.2, 63 x objective, NA 1.40). The images were processed using Fiji software, and the protrusions were quantified using the Neuron Studio program (Rodriguez *et al*, 2008).

### Statistical Analysis

Statistical analyses and graphical representations were generated using GraphPad Prism 10 (version 10.1.2, GraphPad Software, San Diego, CA, USA). For statistical comparison between two groups unpaired-sample t tests were used. Statistical comparison of multiple groups was performed by an analysis of variance (one-way ANOVA) followed by Šídák’s multiple comparisons test for post-hoc comparisons as indicated in figure legends. Collected numerical data are presented in supplementary tables. Data is presented as box with whiskers (25^th^ to 75^th^ percentiles), whiskers represent min to max, the line indicates median and mean is shown as ‘+’.

### LTP recordings in hippocampal slices

Each mouse was euthanized by cervical dislocation, followed by decapitation. The brain was extracted from the skull and transferred into ice-cold ACSF, saturated with carbogen (95% O_2_ / 5% CO_2_) containing (in mM) 250 sucrose, 25.6 NaHCO_3_, 10 glucose, 4.9 KCl, 1.2 KH_2_PO_4_, 2 CaCl_2_, and 2.0 MgSO_4_ (pH 7.3). Both hippocampi were dissected out and sliced transversally (400 µm) using a tissue chopper with a cooled stage (custom-made by LIN, Magdeburg, Germany). The slices were maintained at room temperature in carbogen-bubbled ACSF (95% O_2_ / 5% CO_2_) containing 124 mM NaCl for a minimum of two hours prior to the commencement of the recording. Recordings were performed in the same solution in a submerged chamber that was continuously superfused at 32°C with carbogen-bubbled ACSF (1.2 ml/min). Recordings of field excitatory postsynaptic potentials (fEPSPs) were conducted in the *stratum radiatum* of the CA1b subfield with a glass pipette filled with ACSF. The resistance of the pipette was 1.4 MΩ. The stimulation pulses were applied to the Schaffer collaterals via a monopolar, electrolytically sharpened and lacquer-coated stainless steel electrode, which was positioned approximately 300 μm closer to the CA3 subfield than the recording electrode. Basal synaptic transmission was monitored at a frequency of 0.05 Hz. Long-term potentiation was induced by applying five theta-burst stimulations (TBS) with an interval of 20 seconds. One TBS consisted of a single train of 10 bursts (4 pulses at 100Hz) separated by 200 ms, with a duration of single pulses of 0.2 ms for each pulse. The stimulation strength was set to provide baseline fEPSPs with slopes of approximately 50% of the subthreshold maximum.

The data were recorded at a sampling rate of 10 kHz and then filtered between 0 and 5 kHz. They were subsequently analyzed using IntraCell software, which was developed in-house by LIN Magdeburg in Germany. The statistical evaluation of the mean changes in slope of fEPSPs during the final 10 minutes of recordings was conducted using the Student’s t-test in SigmaPlot (Systat Software Inc., Chicago, IL, USA).

The human recombinant Tissue Inhibitor of Metalloproteinase-3, rhTIMP-3 (R&D Systems, Minneapolis / USA, Cat# 973-TM), was dissolved in distilled water at a concentration of 5 µM and stored at −20°C in small amounts. For each experiment, the final concentration of 15nM was freshly prepared in ACSF and was bath-applied after 10 min stable baseline recording starting 30 min before TBS and ending 10 min after TBS stimulation. For control experiments the appropriate amount of bi-distilled water (vehicle) was applied. The data presented are from 10 slices/5 mice for the control group and 7 slices/4 mice for the TIMP-3 group.

## Acknowledgement

We thank Kathrin Hartung and the team of the animal facility at the LIN Magdeburg for their excellent technical support throughout the project. We thank Eckart Gundelfinger and Helmut Brandstätter for their support and vivid discussions. This project was funded by German Research Foundation (DFG) FR 2758/3–1 and project 460333672 CRC1540 EBM (C02) to RF. Work in the authors CIS and AD labs is supported by the DFG 362321501/RTG 2413 SynAGE and 425899996/CRC1436.

## Author contributions

**Jeet Bahadur Singh**: Conceptualization, Investigation, Methodology, Data curation, Writing original draft, Visualization

**Bartomeu Perelló-Amorós**: Writing-review and editing, Formal analysis, Visualization

**Jenny Schneeberg**: Investigation, Data curation

**Constanze I. Seidenbecher**: Funding acquisition, Resources, Writing-review and editing

**Anna Fejtová**: Funding acquisition, Writing-review and editing, Resources

**Alexander Dityatev**: Conceptualization, Supervision, Funding acquisition, Writing-review and editing, Formal analysis

**Renato Frischknecht**: Conceptualization, Project administration, Supervision, Funding acquisition, Visualization, Formal analysis, Writing original draft, review and editing.

